# Prediction performance of a cardiovascular risk assessment tool using Stanford EHR data repository

**DOI:** 10.1101/648956

**Authors:** Mehrdad Rezaee, Arsia Takeh, Igor Putrenko, Andrea Ganna, Erik Ingelsson

## Abstract

**Background:** Stratification of individuals for their risk to develop cardiovascular diseases can be used for effective prevention and intervention. A significant amount of information for risk assessment can be obtained through repurposing electronic health records (EHR). The objective of this study is to derive and assess the performance of prediction models for cardiovascular outcomes by using EHR-derived data.

**Methods:** We used the Stanford Medicine Research Data Repository (STARR) data from 2000-2017, containing over 2.1 million patients. A subset of 762,372 individuals with complete International Classification of Diseases (ICD) data was used to fit Cox proportional hazard models for prediction of six cardiovascular-related diseases and type 2 diabetes.

**Results:** The derived prediction models indicated consistent high discrimination performance (C-index) for all diseases examined: coronary artery disease (0.85), hypertension (0.82), type 2 diabetes (0.77), stroke (0.76), atrial fibrillation (0.82) and abdominal aortic aneurysm (0.77). Lower prediction abilities were observed for deep vein thrombosis (0.67). These results were consistent across age groups and maintained good prediction abilities among individuals with pre-existing diabetes or hypertension. Assessment of model calibration is ongoing.

**Conclusions:** We proposed new prediction models for the seven diseases using ICD codes derived from EHR data. EHR data can be used for health risk assessment, but challenges related to data quality and model generalizability and calibration remain to be solved.

## Introduction

Cardiovascular diseases (CVD), including diabetes, are the leading causes of global morbidity, mortality and health care costs [1–3]. Appropriate early risk assessment can identify high-risk individuals, spare those at lower risk from intensive interventions, educate patients, provide information on population outcomes, and help in resource allocation [4–6]. In order to accurately determine the risk of developing cardiovascular events, the multifactorial nature of these chronic diseases needs to be considered.

Tools for evaluating cardiovascular risk have been available since the Framingham investigators developed the original algorithms in 1960s [7–10]. Since then, multitudes of other risk assessment approaches have been developed that incorporate an increasing number of risk factors, biomarkers, and comorbidities [10–14]. Clinical application of these models, either for individual patient care or for the purpose of population disease management, requires validation across different patient populations and data sources [15, 16]. There are considerable factors that hinder the performance analysis of any cardiovascular risk model; this is primarily due to the difficulty in obtaining standardized, pertinent large data sets that can be used for calibration and validation studies [17–19].

EHR-based information represents a valuable source of data that can be used for evaluation of prediction modeling. EHRs have the distinct advantage of containing extensive biomedical information on large numbers of individuals across multiple data points. The purpose of the current study is to use EHR information from a large population of patients (STARR) to evaluate the performance of newly developed risk assessment models using International Classification of Diseases (ICD) outcomes derived from the EHR data.

## Material and Methods

### Data source

We obtained de-identified patient data from the STARR dataset. The STARR population (2.1 million) includes patients from all ages (0 to 91 years old) who have attended Stanford Hospital or any of its clinics from 2001 to 2017. Demographic information, encounters, lab results and pharmacy orders are recorded in the database. However, conventional cardiovascular risk factors (biomarkers, diabetes status and family history) had a high frequency of missing data (80-100 percent). For this reason (i.e. the complete data was limited) we looked into alternative types of data. Specifically, we focused on using only ICD-derived (International Classification of Diseases, Tenth Revision, Clinical Modification) risk factors for prediction of the cardiovascular-related events. Thus, we included all individuals who were 18 years-old or older, who had at least one ICD code for the relevant cardiovascular risk factors and had non-missing information on age, BMI and sex. This decreased the population size to 762,372.

### Risk factors and outcome definitions

In addition to age, BMI and sex, the presence of cardiovascular-related risk factors was defined using the following ICD-10 codes: elevated triglycerides (E78.1), elevated cholesterol (E78.00), depressed HDL cholesterol (E78.6), elevated LDL Cholesterol (E78.5), elevated creatinine (R79.89) and vitamin D deficiency (E55.9).

Disease outcomes were defined based on the following ICD codes: coronary artery disease (CAD): I20–I25 and T82 codes; hypertension (HTN): I10, I15, and R03.0 codes; type 2 diabetes mellitus (DM); E11, E13, and E14 codes, stroke: G46.3, G46.4, I63, I66, I67, and I693 codes; deep vein Thrombosis (DVT): H34.8, H40.8, I23.6, I24.0, I63, I67.6, I74, I81, I82, I87.2, I87.3, K64.5, N48.8, N52.0, O03.3, O03.8, O04.8, O07.3, O08.7, 022, O87, Q26, T82.8, T83.8, T84.8, T85.8, and Z86.7 codes; abdominal aortic aneurysm (AAA): I71 and I79.0 codes; and for atrial fibrillation (AF): I48-49 codes. For each disease endpoint, we also obtained the date when the patient was diagnosed with the endpoint.

### Statistical Analysis

We consider risk factors recorded from 2001 to 2012 and use these risk factors to predict five-year risk for each disease outcome between 2012 and 2017. For each disease outcome, we also excluded individuals who were diagnosed with that disease before the baseline year of 2012. That is, we considered only incident outcomes. End-of-follow up is defined as the diagnosis of the disease, death or end of the study (December 2017).

Linear Cox proportional hazard (PH) models were developed using lifelines 0.13.0 Python library.

Since the risk factors were not measured at baseline, but during an 11 year period, they were modeled as time-varying covariates. If multiple instances of the same risk factors were measured for the same individual we considered the last instance, as this is the closest to the baseline.

The discrimination was assessed on the basis of five-fold cross-validated Harrell’s concordance index (C-index) [20–22]. The cross-validated C-index was used as a main metric for assessing discriminative ability of the Cox PH-based models.

Principal component analysis (PCA) was used for validating the selection of variables and to avoid overfitting through comparison of the number of selected variables and optimal number of principal components [23]. The number of components to be retained was determined by using maximum-likelihood density estimation and full singular value decomposition as parameters of the PCA function, which applies Bayesian model selection to probabilistic PCA in this configuration.

### Sensitivity Analysis

To evaluate the prediction performances of the derived models across different subsets of the population we performed sensitivity analyses in the following subpopulations: 1) healthy participants without any of the cardiovascular-related diseases or type 2 diabetes at the baseline, 2) the following age groups, with age measured at baseline: <45, 45-55, 56-65, 66-75, >75, 3) individuals with at least one non-target disease (DM, HTN, Stroke, DVT).

## Results

The study population included 762,372 adult patients ages 18 years old and above, which visited Stanford Hospital and clinics during the period spanning 2000-2017, had at least one cardiovascular-related ICD code and information on age, sex and BMI (Table 1). We notice that this approach reduces the generalizability of our results. Nonetheless, this study represents a proof of concept that prediction models can be derived from ICD data. Table 2 reports the number of individuals included in each disease-specific analysis and the incidence of the main disease outcome for general population and age stratified sub-populations. Individuals reported in Table 2 were used to derive the final predictions. As presented in Table 3, the cross-validated discrimination metrics (C-index) across all disease was high (>0.75) except for DVT with a C-index of 0.67.

**Table 1:** Characteristics of the population included for the final analysis (762, 372). Number of individuals in each group, on brackets, percentage of individuals in each group.

| Gender | No. [%] |
| --- | --- |
| Male | 412,062 [54.05] |
| Female | 350,310 [45.94] |
| Race |  |
| White | 368,378 [48.32] |
| African-American | 28,436 [3.73] |
| Asian | 91,103 [11.95] |
| Unknown | 142,182 [18.26] |
| Other | 135,244 [17.74] |
| Age |  |
| 18-35 | 182,969 [23.99] |
| 35-45 | 126,554 [16.60] |
| 45-55 | 134,940 [17.70] |
| 55-65 | 129,603 [17.00] |
| 65-75 | 109,019 [14.30] |
| 75+ | 79,287 [10.40] |

**Table 2:** Population size and disease incidence. Incidence (number of incident cases/total population) for each target disease is presented for the total population as well as age groups for each disease. Numbers inside the parenthesis are the Incidence percentages.

|  | CAD | DM | HTN | Stroke | AF | DVT | AAA |
| --- | --- | --- | --- | --- | --- | --- | --- |
| General | 25,409 / | 30,652 / | 101,901 / | 8,810 / | 23,035 / | 13,857/ | 3,260 / |
|  | 762,372 | 762,372 | 762,372 | 762,372 | 762,372 | 762,372 | 762,372 |
|  | (3.33%) | (4.02%) | (13.37%) | (1.16%) | (3.02%) | (1.82%) | (0.43%) |
| < 45 | 1202 / | 4,504 / | 14,803 / | 1,046 / | 1,421 / | 2,301 / | 322 / |
|  | 309,523 | 309,523 | 309,523 | 309,523 | 309,523 | 309,523 | 309,523 |
|  | (0.39%) | (1.46%) | (4.78%) | (0.34%) | (0.46%) | (0.74%) | (0.10%) |
| 46-55 | 2,631 / | 5,384 / | 17,312 / | 1,039 / | 1,970 / | 2,253 / | 337 / |
|  | 134,940 | 134,940 | 134,940 | 134,940 | 134,940 | 134,940 | 134,940 |
|  | (1.95%) | (3.99%) | (12.83%) | (0.77%) | (1.46%) | (1.67%) | (0.25%) |
| 56-65 | 5,611 / | 7,724 / | 23,432 / | 1,620 / | 4,070 / | 2,955 / | 609 / |
|  | 129,603 | 129,603 | 129,603 | 129,603 | 129,603 | 129,603 | 129,603 |
|  | (4.33%) | (5.96%) | (18.08%) | (1.25%) | (3.14%) | (2.28%) | (0.47%) |
| 66-75 | 8,100 / | 7,958 / | 25,129 / | 2235 / | 6,552 / | 3,216 / | 1,025 / |
|  | 109,019 | 109,019 | 109,019 | 109,019 | 109,019 | 109,019 | 109,019 |
|  | (7.43%) | (7.30%) | (23.05%) | (2.05%) | (6.01%) | (2.95%) | (0.94%) |
| > 75 | 7,865 / | 5,082 / | 21,225 / | 2,870 / | 9,022 / | 3,132 / | 967 / |
|  | 79,287 | 79,287 | 79,287 | 79,287 | 79,287 | 79,287 | 79,287 |
|  | (9.92%) | (6.41%) | (26.77%) | (3.62%) | (11.38%) | (3.59%) | (1.22%) |

**Table 3:** Five-fold cross validated C-index [95% CI] as a measure of prediction performance for each disease.

| Disease | General |
| --- | --- |
| CAD | 0.86 [0.83-0.90] |
| DM | 0.77 [0.75-80] |
| HTN | 0.82 [0.79-0.86] |
| Stroke | 0.76 [0.72-0.79] |
| AF | 0.82 [0.80-0.85] |
| DVT | 0.67 [0.64-0.70] |
| AAA | 0.77 [0.73-0.80] |

Modeling performance was further evaluated across clinically-distinct sub-populations (Table 4). Prediction models for each disease had consistent behavior across all the subpopulations examined.

**Table 4:**
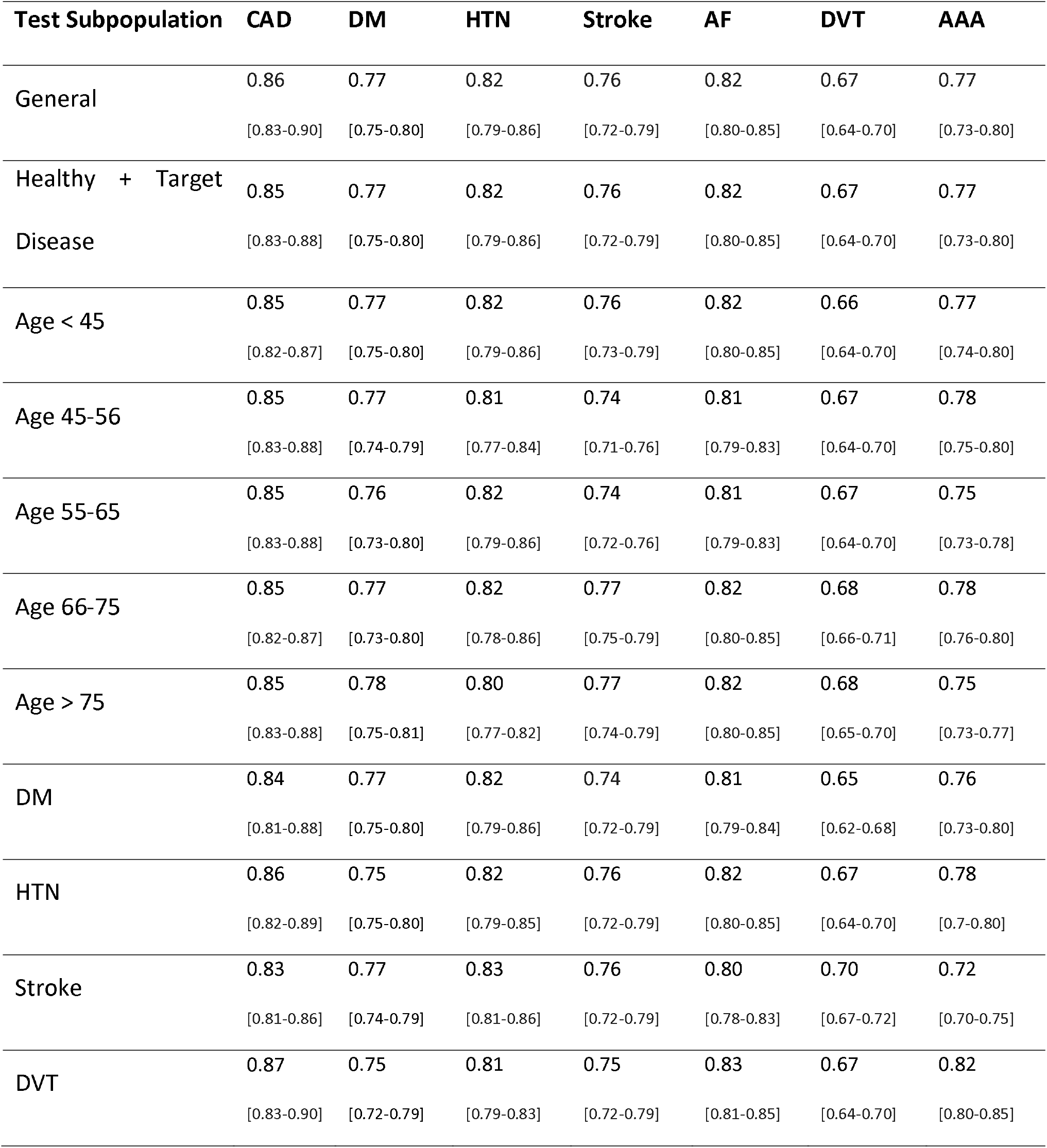
Discriminative ability of risk prediction models across different subgroups.

| Test Subpopulation | CAD | DM | HTN | Stroke | AF | DVT | AAA |
| --- | --- | --- | --- | --- | --- | --- | --- |
| General | 0.86<br>[0.83-0.90] | 0.77<br>[0.75-0.80] | 0.82<br>[0.79-0.86] | 0.76<br>[0.72-0.79] | 0.82<br>[0.80-0.85] | 0.67<br>[0.64-0.70] | 0.77<br>[0.73-0.80] |
| Healthy + Target Disease | 0.85<br>[0.83-0.88] | 0.77<br>[0.75-0.80] | 0.82<br>[0.79-0.86] | 0.76<br>[0.72-0.79] | 0.82<br>[0.80-0.85] | 0.67<br>[0.64-0.70] | 0.77<br>[0.73-0.80] |
| Age < 45 | 0.85<br>[0.82-0.87] | 0.77<br>[0.75-0.80] | 0.82<br>[0.79-0.86] | 0.76<br>[0.73-0.79] | 0.82<br>[0.80-0.85] | 0.66<br>[0.64-0.70] | 0.77<br>[0.74-0.80] |
| Age 45-56 | 0.85<br>[0.83-0.88] | 0.77<br>[0.74-0.79] | 0.81<br>[0.77-0.84] | 0.74<br>[0.71-0.76] | 0.81<br>[0.79-0.83] | 0.67<br>[0.64-0.70] | 0.78<br>[0.75-0.80] |
| Age 55-65 | 0.85<br>[0.83-0.88] | 0.76<br>[0.73-0.80] | 0.82<br>[0.79-0.86] | 0.74<br>[0.72-0.76] | 0.81<br>[0.79-0.83] | 0.67<br>[0.64-0.70] | 0.75<br>[0.73-0.78] |
| Age 66-75 | 0.85<br>[0.82-0.87] | 0.77<br>[0.73-0.80] | 0.82<br>[0.78-0.86] | 0.77<br>[0.75-0.79] | 0.82<br>[0.80-0.85] | 0.68<br>[0.66-0.71] | 0.78<br>[0.76-0.80] |
| Age > 75 | 0.85<br>[0.83-0.88] | 0.78<br>[0.75-0.81] | 0.80<br>[0.77-0.82] | 0.77<br>[0.74-0.79] | 0.82<br>[0.80-0.85] | 0.68<br>[0.65-0.70] | 0.75<br>[0.73-0.77] |
| DM | 0.84<br>[0.81-0.88] | 0.77<br>[0.75-0.80] | 0.82<br>[0.79-0.86] | 0.74<br>[0.72-0.79] | 0.81<br>[0.79-0.84] | 0.65<br>[0.62-0.68] | 0.76<br>[0.73-0.80] |
| HTN | 0.86<br>[0.82-0.89] | 0.75<br>[0.75-0.80] | 0.82<br>[0.79-0.85] | 0.76<br>[0.72-0.79] | 0.82<br>[0.80-0.85] | 0.67<br>[0.64-0.70] | 0.78<br>[0.7-0.80] |
| Stroke | 0.83<br>[0.81-0.86] | 0.77<br>[0.74-0.79] | 0.83<br>[0.81-0.86] | 0.76<br>[0.72-0.79] | 0.80<br>[0.78-0.83] | 0.70<br>[0.67-0.72] | 0.72<br>[0.70-0.75] |
| DVT | 0.87<br>[0.83-0.90] | 0.75<br>[0.72-0.79] | 0.81<br>[0.79-0.83] | 0.75<br>[0.72-0.79] | 0.83<br>[0.81-0.85] | 0.67<br>[0.64-0.70] | 0.82<br>[0.80-0.85] |

## Discussion

The extensive availability of EHR data offers a unique and promising opportunity for the application of prediction models at the point of care or for population planning. EHR data have the advantage of including large numbers of individuals and recording many interactions that an individual has with the health system. This aspect allows us to model longitudinal changes in risk factors. The main disadvantage of the EHR-based data is low data quality, poor generalizability outside the specific EHR system and loss at follow-up (i.e. inability to track people that change health systems or emigrate).

The models derived in this paper focus on the same set of diseases that have been the focus of a companion paper developed on a large epidemiological cohort study (UK Biobank) [24]. Despite different data sources, we observed comparable prediction abilities across the studies. However, one advantage of large epidemiological studies as compared to EHR-based data is the limited number of missing data that allows for direct modeling of biomarkers as risk factors. In the current study this was not possible, and we had to rely on ICD-based diagnosis. This is suboptimal as the lack of ICD codes does not necessarily imply that the risk factor is not present in the individual. Moreover, this binarizes the underlying continuous variables (e.g. cholesterol), which limits the predictive value of such risk factors. Nonetheless, we still observe good prediction abilities indicating that the current, pragmatic approach has some value.

In this study we didn’t evaluate model calibration. This is a limitation that will be addressed in a future revision of the manuscript.

## Conclusions

In this report, we present the development and validation of risk prediction models for cardiovascular-related diseases and type 2 diabetes using EHR data. Due to the large number of missing data for traditional risk factors, ICD codes were used to define the model predictors. The developed models had good prediction abilities in the entire study population as well in specific clinically-relevant subgroup populations. Future research will focus on including additional risk factors, such as biomarkers and genetic information, and on evaluating the generalizability outside this specific study population. Prediction models derived from EHR data have the potential to be used for primary prevention of cardiovascular-related disease.

## Acknowledgement

Precision wellness, Inc. was solely responsible for the conception and development of the risk analysis algorithms described in this report.

EI served as the PI for this study, which was performed, in part, through a sponsored research (SPO 134382) agreement with Stanford University.

## Funding Source

The funder, Precision Wellness, Inc., provided support in the form of salaries for authors AT and IP, consultancy fees to AG, and as an unrestricted research grant to Stanford University (led by EI), but did not have any additional role in the study design, data collection and analysis, decision to publish, or preparation of the manuscript. The specific roles of these authors are articulated in the ‘Author contributions’ section.

## Author Contributions

1. **Conceptualization:** MR AT
2. **Data curation:** AT
3. **Formal analysis:** AT, IP
4. **Funding acquisition:** MR
5. **Investigation:** MR AT EI
6. **Methodology:** MR AT IP AG EI
7. **Project administration:** MR AT
8. **Resources:** MR
9. **Software:** AT
10. **Supervision:** MR AT
11. **Validation:** MR EI AG
12. **Visualization:** AT, IP
13. **Writing – original draft:** AT IP MR
14. **Writing – review & editing:** MR EI AG

